# Pathogen-derived mechanical cues regulate the spatio-temporal implementation of plant defense

**DOI:** 10.1101/2021.10.18.464859

**Authors:** Ophélie Léger, Frédérick Garcia, Mehdi Khafif, Nathalie Leblanc-Fournier, Aroune Duclos, Vincent Tournat, Eric Badel, Marie Didelon, Aurélie Le Ru, Sylvain Raffaele, Adelin Barbacci

## Abstract

How immune responses are activated and regulated is a central question in immunology. In addition to molecular signaling, recent work has shown that physical forces regulate the immune response of vertebrates by modifying transmembrane protein conformation and cell contact. Mechanical stress and strain produced by forces constitute physical cues perceived by cells instructing gene expression. Whether mechanical cues generated by pathogens during host colonization can trigger adaptive responses in plant cells remains elusive. We found that local and progressive variations of plant cell wall tension caused by fungal pathogen attacks are transmitted to neighboring healthy tissue around the infection site and trigger immunity in distal cells. This ‘thigmoimmunity’ process requires the reorganization of cortical microtubules and contributes strongly to Arabidopsis disease resistance.

**One-Sentence Summary:** Activation of plants immunity depends on fluctuations of mechanical tension caused by a fungal pathogen.

## Main Text

Plants are composed of fixed and bound cells that form a solid-like continuous material favorable to the spread of mechanical cues. Plant tissues are under high internal mechanical stress (homogeneous to a force per surface unit) generated by the interaction of water in the vacuole and the stiff polysaccharides wall. Tensile stresses in the cell wall serve as cues for cells to read their own shapes. The monitoring of the stress field in cell walls is a central mechanism to the plant perception of self, coined proprioception (**1, 2**). Many plant pathogens derive carbon from cell walls of their host for growth and reproduction. Paradigmatic examples are necrotrophic fungal pathogens that secrete enzymes and organic acids to exploit carbon sources in plant cell walls (**3**). Current models of plant immunity involve the perception of microbial molecules, free carbohydrates and other host degradation products for the activation of localized defense (**4**). The transfer of signal molecules is proposed to inform rapidly distant cells of the occurrence and position of a pathogen attack triggering systemic immunity (**5**). Signals governing plant systemic immunity remain however incompletely understood (**6**). By challenging plant cell wall integrity, pathogens affect mechanical stress constantly monitored by cells for proprioception (**1, 2**). Because plant tissues are solid like continuous structures, pathogen-derived mechanical stresses that induces shape variation described by strain, are not local only and could reshuffle internal stress across cells in an quasi-instantaneous and oriented manner and inform distant cells. Alterations to stress in plant cell walls result in the formation of measurable mechanical strain underpinning variations in cell shape. However, whether plant cells directly sense and respond to mechanical stimuli induced by pathogen colonization has not been formally established yet. This prompted us to ask whether such mechanoperception of pathogen-derived stress and strain could act as a signal controlling immune responses in plants.

### Pathogen-derived mechanical cues guide microtubules organization

To analyze the mechanical consequences of infection by the necrotrophic fungal pathogen *Sclerotinia sclerotiorum*, we monitored the 2D kinematics of *Arabidopsis* deformation in epidermal cells expressing the microtubule-associated protein 4 (MAP4) fused to GFP over two hours time course (24hpi to 26hpi). Since reshuffling of mechanical stress in walls leads to cells deformations, we combined time-lapse confocal imaging and an optical flow tracking method (**7**) to quantify cell shape variation. We found that infection by *S. sclerotiorum* generated excessive deformations in healthy cells next to the inoculation site, due to the global reorganization of mechanical stress balance in walls (**Fig.1A**). Relative surface shape variations were extracted from the strain tensor and computed as the first invariant. The mean magnitude of variations was on absolute value ca. 1% of the initial surface and twice higher in infected than in healthy tissues (**Fig.1C**). Deformations affected all observed cells and were not limited to cells directly in contact with the pathogen (**Fig.1A**). Cell surfaces were either shrunken or stretched, suggesting that the mechanical stress reshuffling necessary to reach the mechanical equilibrium resulted from the balance between local shape-derived stress and pathogen-derived stress. The pathogen-derived strain pattern was weakly tangential to the periphery of the area colonized by the fungus (**Fig.1A**), in contrast with the circumferential stress pattern typically observed after laser ablation of cells (**8,9**). To explore possible causes for this discrepancy, we derived a 2D mechanical model of leaf tissue at a similar scale to confocal observations in which we mimicked the mechanical effect of pathogen inoculation (**Fig.1B**). The model showed that a heterogeneous release of the initial isotropic tension leads to a non-circumferential strain pattern around the infection site (**Fig.1B**). We conclude that the non-circumferential stress patterns we observed are due to the heterogeneous release of mechanical stress in leaf tissue, consistent with *S. sclerotiorum* causing partial cell wall loosening through the chemical activity of enzymes and organic acids (**3**).

**Fig. 1.**
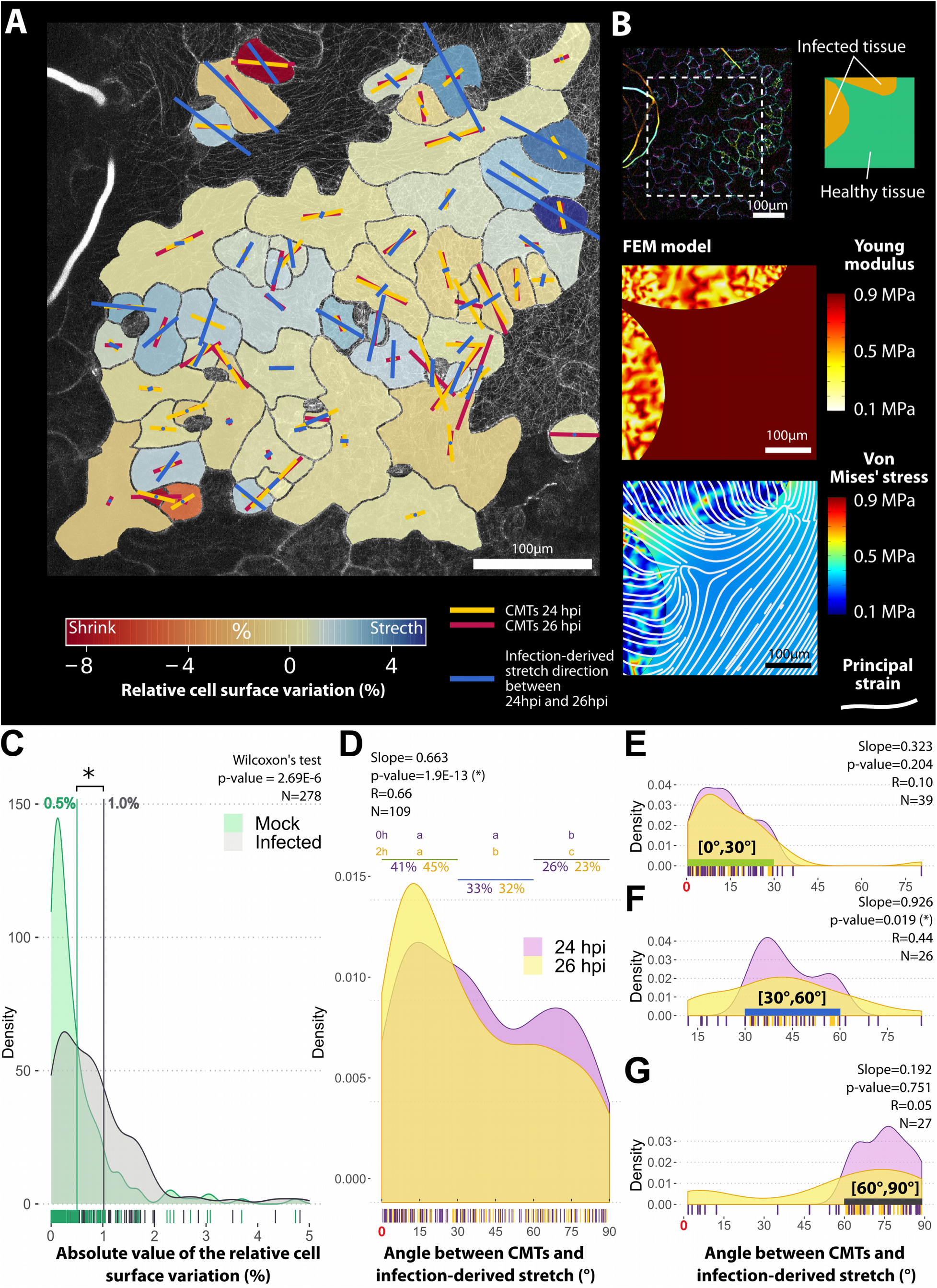
Pathogen-derived stretch guides the reorganization of cortical microtubules (CMTs). (**A**) Experimental analysis of pathogen-derived stretch (blue segments), temporal orientation of the CMT network at 24 hours post inoculation (hpi,yellow segments) and 26 hpi (red segments) in *A. thaliana* leaf epidermal cells. Segments length codes for the CMT anisotropy. Cell surface variation is indicated as a colored background. (**B**) Mechanical finite element model (FEM) of infection-derived strain. Around infected tissue (left up side of the region of interest), the partial release of internal stress and Young’s modulus led to non-circumferential strain patterning (**C**) Distribution of the magnitude of the relative cell surface variation triggered by pathogen-derived stress release. Individual measures are shown as bars along the X-axis, the average was 1% in infected tissue and 0.5% healthy tissue. (**D**) Distribution of CMTs angle relative to the pathogen-derived stretch direction at 24 hpi and 26 hpi. The proportion of cells with CMTs oriented within 0-30°, 30-60° and 60-90° of the pathogen-derived stretch is indicated above the graph, and compared at 24 and 26hpi using a proportion test. Distribution of CMTs angle relative to the pathogen-derived stretch at 24 and 26hpi for cells with CMTs within 0-30° (**E**), 30-60° (**F**) and 60-90° (**G**) of the pathogen-derived stretch. The slope of the linear model indicated the temporal trends of angle evolution over 2h. Slope below 1 indicates a global reorganization of CMTs in the direction of the infection-derived stretch.

Laser ablation experiments showed that mechanoperception occurs when the stress release exceeds shape-derived stress (**9**). In such case, mechanoperception is associated with the anisotropic reorganization of the cortical microtubules (CMTs) in the direction of the tensile bias (**9, 10**). Consistent with these results, cortical microtubules in infected tissue were more anisotropically organized than in healthy tissues (**Fig.S1**). The signal instructing the reorientation of CMTs must therefore encode the direction of the bias, i.e. encode 3D information (described by a tensorial field). While diffusive molecules more likely encode 1D information (described by a scalar field), mechanical stress and strain, 3D by nature, are prime signal candidates instructing CMTs reorganization (**10**).

In the absence of growth (plastic deformation), *in vitro* and *in vivo* experiments suggest that CMTs align in the direction of the stretch, i.e. the elastic elongation (**10**). To determine whether cells perceive pathogen-derived stress, we monitored the angle between pathogen-derived stretch and CMTs in *Arabidopsis* epidermal cells over two hours (24 to 26 hpi). At 24 hpi, a majority of cells exhibited CMTs already oriented between 0° and 60° from the stretch direction (**Fig.1D**). Nevertheless, we detected a significant reorganization of CMTs parallel to the stretch during the subsequent two hours. During this period, the proportion of CMTs within 30° of the stretch direction increased from 41% to 45% (**Fig.1D**). When closely aligned or orthogonal with stretch, CMTs orientation remained stable (**Fig1.E,G**). Alignment of CMTs along the stretch was mostly due to the reorganization of CMTs initially oriented from 30° to 60° of the stretch (**Fig.1F**). Altogether, these results suggested that plant cells perceived pathogen-derived tensile stretches as mechanical cues guiding the reorganization of CMTs.

### Pathogen-derived mechanical cues are not cell-autonomous

Plant tissue structure favored potentially the nearly instantaneous propagation of mechanical stress and strain that may inform distant cells of the presence and the position of a pathogen. Our observations suggested that the perception of pathogen-induced mechanical stress was not restricted to cells immediately adjacent to the colonized area. To estimate the distance to which mechanical cues reshuffle internal stress during infection, we designed a 2D mechanical model at the scale of the leaf to compute the spatial stress release caused by infection (**Fig.2A**). The progress of infection over time was modeled by the radial growth of a virtual pathogen colony from 0.5 to 10 mm in a leaf modeled. Simulations showed that the pathogen-derived stress developed an overstretch ring adjacent to the infection site (**Fig.2A,B**). The width of the overstretched ring depended only on the lesion radius (**Fig.2B**). For pathogen colonies smaller than 6 mm radius, the width of the overstretch ring was greater than 2 mm (Fig.2B). Importantly, the model supports a spread of the mechanical signal independently of signaling molecules. In agreement with the simulations, we observed the reorganization of CMTs over 2 hours (24-26 hpi) in a 2.5mm-wide ring around hyphae of *S. sclerotiorum* in *Arabidopsis* leaf epidermal cells expressing the MAP4:GFP construct. At 24 hpi, 43% of the CMTs within ca. 0.5 mm from the fungal hyphae were within 30° of the direction of the infection-derived stretch and remained stable over 2 hours (**Fig.2C**). At c.a. 2.5mm from the infection site, the proportion of CMTs within 30° of the direction of pathogen-derived stretch increased from 31% to 38% (**Fig.2C**). The global organization of CMTs along the direction of the stretch appeared dependent on the distance from the infection site. Close to the infection site, the alignment of CMTs with mechanical cues was already clear at 24hpi, whereas the reorganization was initiated but incomplete in distant cells. We conclude that mechanical stretch by pathogen infection was perceived at a supra-cellular level through a mechanism involving CMTs re-orientation.

**Fig. 2.**
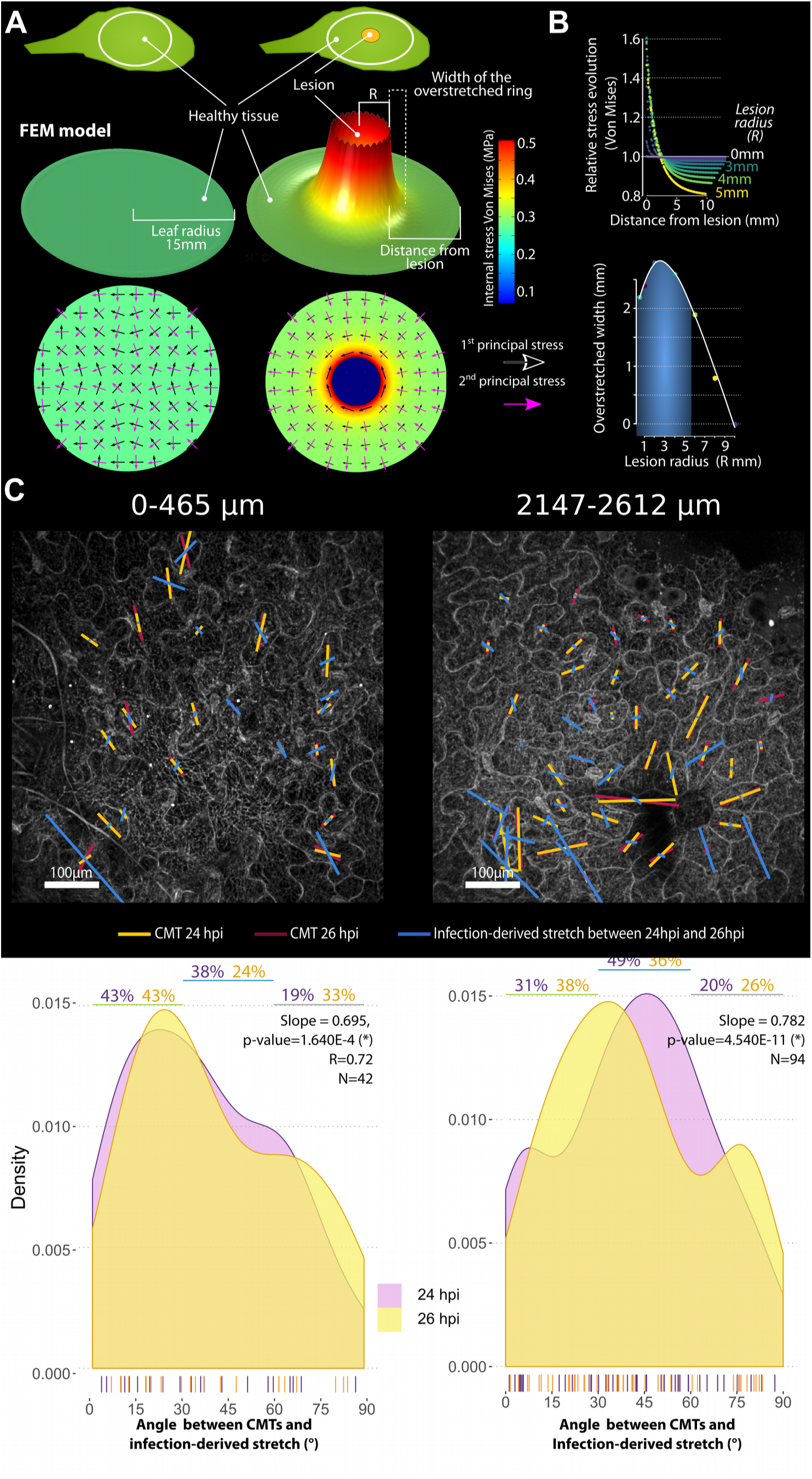
Pathogen-derived stretches shape the reorganization of CMTs in a 2.5 mm ring around the colonized area. (**A**) Biomechanical model of infected leaf. Healthy tissue was initially under tension (black and purple arrows represent principal stresses). The infection releases progressively the internal tension that reshuffle internal stress through tissue and creates a pathogen-derived overstretched ring around the colony. (**B**) Profile of infection-derived stress depended only on the size of the virtual colony. For lesion below 6 mm of radius, the width of the overstretched ring was larger than 2mm. (**C**) Patterns of pathogen-derived stretch and CMT orientation within a 0.5 mm and 2.5mm range from the colonized area at 24 hpi and 26 hpi. CMTs direction tends towards the direction of the infection derived stretch over 2h. Individual measures are shown as bars along the x-axis, the % of cells with CMTs oriented within 0-30°, 30-60° and 60-90° of the pathogen-derived stretch is indicated above the graph, slope of the linear model indicated the temporal trends of the distribution. Slope below 1 indicates a global reorganization of CMTs in the direction of the infection-derived stretch.

### Infection-derived mechanical cues trigger differential genes expression

We reasoned that, if adaptive, the perception of pathogen-induced CMTs reorganization is likely to influence plant immune responses. To test this hypothesis, we took advantage of the *A. thaliana pfd6-1* mutant in Col-0 background altered in the prefoldin 6 gene (At1g29990) leading to defects in CMT dynamics (**11**). First, we measured by quantitative RT-PCR the expression of marker genes associated with mechanoperception and disease resistance in the *pfd6-1* and Col-0 wild type genotypes for healthy and infected plants (**Fig.3**). Samples were extracted from the 2.5 mm overstretched ring around the inoculated or mock-treated area, in which no disease symptoms were visible. During infection by *S. sclerotiorum*, the *AtMCA1* (MID1-COMPLEMENTING ACTIVITY 1) gene expression was downregulated by a similar ratio in Col-0 and *pfd6-1* (**Fig.3A**). *AtMCA1* is involved in mechanoperception (**12**) and cell wall integrity (CWI) maintenance by promoting Jasmonic acid accumulation (**4**). CWI maintenance monitors local and cell-autonomous signals such as the alteration of cell wall components (**4**). Therefore, CWI-triggered defenses may be weakened by *S. sclerotiorum* independently on the CMTs dynamics. The defect in CMTs dynamics did not alter the expression of the gene *AtPRT* (Pathogenesis-related thaumatin superfamily protein) associated with resistance (**Fig.3A**). By contrast, the expression of *AtBON1, AtBON2* and *AtPDR12*, involved in resistance to several fungal pathogens (**13, 14**) was dependent on CMTs dynamics. *AtBON1* and *AtBON2* are Ca2+ dependent copine-like genes expressed at the same level in Col-0 and *pfd6-1* but upregulated in the periphery of infection only in wild type plants (**Fig.3B**). CMTs dynamics also affected the expression of *AtPDR12*, an ABC transporter involved in ABA transport and resistance to fungal pathogens (**15, 16**). While *AtPDR12* was upregulated more than 3500-fold upon *S. sclerotiorum* inoculation in wild type plants, upregulation was only 2.4 fold in *pfd6-1* (**Fig.3B**). Finally, we compared resistance to *S. sclerotiorum* in Col-0 / *pfd6-1* and *WS-4 / bot1-7* plants using image-based time-resolved phenotyping (**17**). *bot1-7* is a katanin mutant allele in the Ws-4 background affected in the CMTs severing process (**9**). The fungal pathogen colonized *pfd6-1* and *bot1-7* significantly faster than their respective wild type, indicating that CMT dynamics contributes at least to ca. 40% of the disease resistance phenotype (**Fig.3B**). Altogether our results show that the mechanoperception of pathogen-derived cues in leaves reshuffle internal stress across a few millimeters range around the infection site and controls the expression of disease resistance genes in distal cells. This mechanical regulation is potentially independent of CWI-triggered defense as currently defined.

**Fig. 3.**
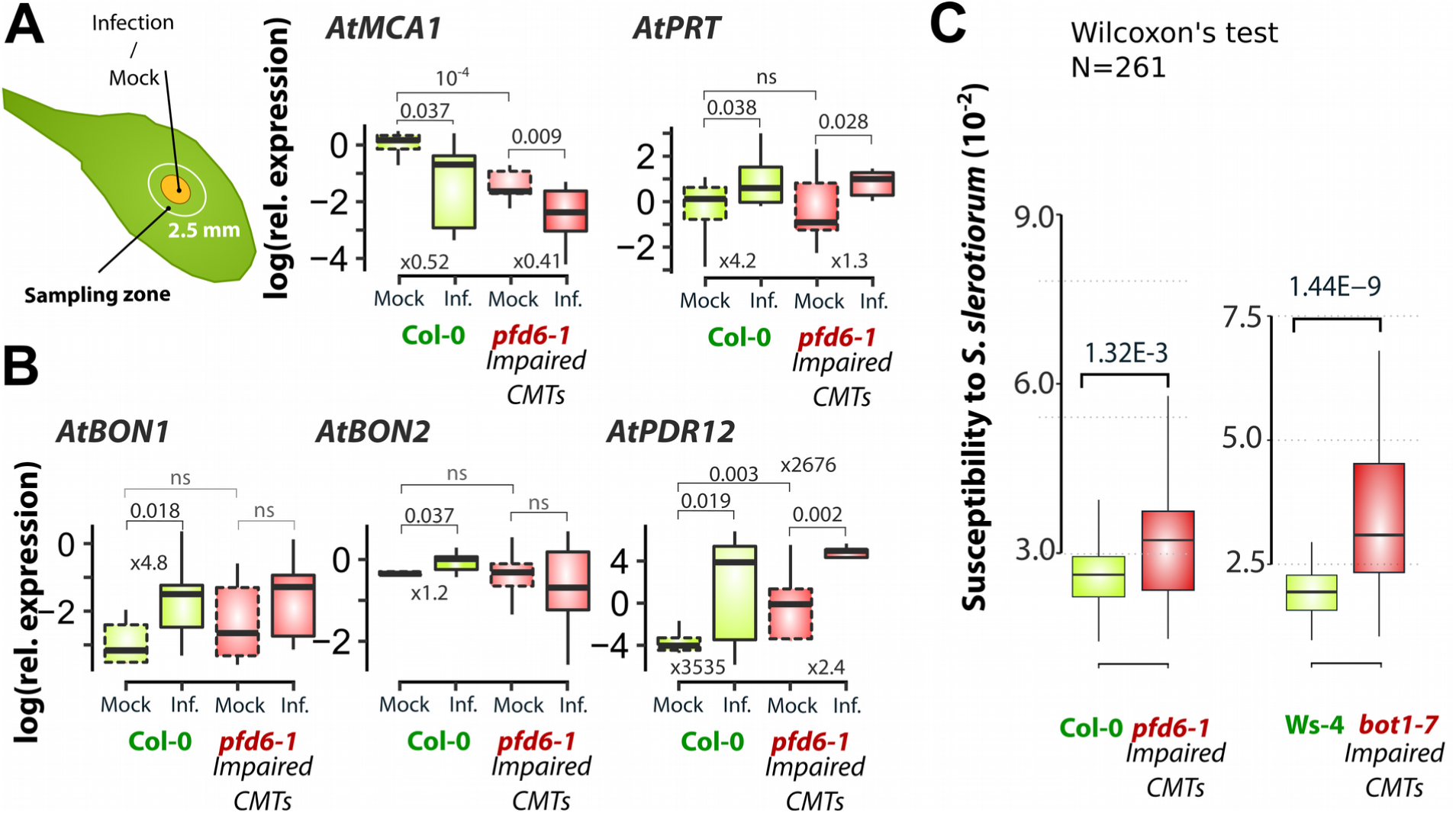
Effect of impaired CMTs dynamics on plant disease resistance. (**A,B**) Relative expression of plant marker genes in cells from the overstretched area surrounding disease lesions determined by quantitative RT-PCR in Col-0 (green) and the *pfd6-1* mutant impaired in cortical microtubules (CMTs) dynamics (red). *AtMCA1*, associated with cell wall integrity perception, and *AtPRT*, associated with immunity, were regulated similarly upon infection (Inf.) in the Col-0 and *pfd6-1* genotypes. By contrast, *AtBON1, AtBON2* and *AtPDR12* were strongly upregulated upon infection in Col-0 but not in *pfd6-1*. Values shown for three independent biological replicates, each analyzed in three technical replicates. Multiple pairwise t-test p-values are indicated over brackets. (**C**) Plant susceptibility to *S. sclerotiorum* fungal pathogen was drastically increased in mutants impaired in CMTs dynamics. Values shown for 261 leaves per genotype, analyzed in three independent assays. Means were compared by Wilcoxon’s test and significant p-value are indicated over brackets. Boxplots show first and third quartiles (box), median (thick line), and the most dispersed values within 1.5 times the interquartile range (whiskers). ns. non significant.

### Concluding remarks

Thigmomorphogenesis describes the connection between mechanoperception and shape regulation (**18**). We describe here a role for CMTs in connecting mechanoperception and defense regulation in plants, a process we propose to refer to as thigmoimmunity. Recent work demonstrated the central role of proprioception in the regulation of shape and posture by monitoring mechanical stress fluctuation in cell walls (**1, 2**). Our findings suggest that proprioception also instructs the plant immune response. The solid-like mechanical behavior of plant tissue favors the propagation of mechanical cues released by infection through the apoplast. Mechanoperception occurs in every cell around the infection and does not require the loss of wall integrity but only the tensorial variation of stress (**19**). Proprioception via mechanical cues also allows the integration of external mechanical stimuli such as those caused by wounding or wind (**1, 2**), implying that the level of plant resistance depends on external mechanical stimulation. Such corollary is in line with the gain of resistance provided by soft external mechanical stimulation observed in *A. thaliana* infected by the necrotrophic fungus *Botrytis cinerea* (**20, 21**).

Our results highlight the mechanoperception of pathogen-derived stress as a distinct layer of the plant defense system with unique properties. Well-established mechanisms of plant immunity are the pathogen-associated molecular patterns (PAMPs)-triggered immunity (PTI) and effector-triggered immunity (ETI) which involve respectively, the perception pathogen-derived molecule and the perception of damage-associated molecular patterns (DAMPs) derived from plant molecules (**22**). The thigmoimmunity layer of plant immunity differs from previously described mechanisms by the non-molecular nature of the signals that activate it. Infection-derived mechanical signals are mobile since local cell wall hydrolysis involves a global stress reshuffling in the vicinity of pathogen-colonized cells. Infection-derived mechanical signals contain oriented 3D information (tensorial), and do not involve molecular diffusion or transport processes. Because of the intrinsic properties of mechanical cues, thigmoimmunity could orchestrate spatially, around the infection site, a non-specific form of resistance and boost molecular immune responses. The spatial propagation of pathogen-derived mechanical cues would explain partly the spatial heterogeneity of gene expression observed during infection (**23**). Recent results suggests that perception of cell damage caused by infection is not mediated by DAMPs only and involves additional cues (**6**). We propose that pathogen-derived stretch, generated across plant cells by the local release of internal tension, acts as additional non-molecular cues triggering defense response to fungal infection. While mechanical cues can encode spatial information, they are unspecific and cannot inform cells of the nature of the attacking pathogen, in contrast with molecular-based defense layers. The thigmoimmunity layer complement thus the signals involved in PTI and ETI. Altogether, layers of plant defense orchestrate a complex spatio-temporal response to pathogens.

## Materials and Methods

### Plant / Fungus materials, culture conditions, and inoculation

*Arabidopsis thaliana* plants were grown at 23°C with 9 hours/day of light for 5 weeks. *Sclerotinia sclerotiorum* was grown on PDA (Potato Dextrose Agar, Fluka) plates at 23°C for 4 days in the dark prior to inoculation. Inoculations were performed by depositing a 5mm diameter plug of culture medium containing mycelium on the upper surface of leaves. Mock treatment consisted in a 5mm diameter plug of mycelium-free PDA. For quantitative RT-PCR analyses, inoculated plants were placed in mini-greenhouses during 24 hours to keep a high humidity level promoting infections. For phenotypic analysis, 3 leaves per plant were cut and put in Navautron devices to monitor the kinematic of lesion development (**24**).

### Quantitative RNA analysis

Plant samples used for the measurement of gene expression were collected 24 hours post inoculation (hpi). The leaves were cut at the base of the petiole and immediately placed on a glass plate cooled by liquid nitrogen. For each leaf, 2.5 mm wide leaf rings were cut around the necrosis and stored at −80°C. RNA extraction was performed using the Nucleospin RNA kit (Macherey-Nagel). cDNA was synthesized with Superscript III reverse transcriptase (Invitrogen Carlsbad, CA) using 1 μg of total RNA. Quantitative RT-PCR was performed using gene-specific primers (**Table S1)**, on a Light Cycler 480 II apparatus (Roche Diagnostics, Meylan, France) using Roche SYBR green I reagent. Five pmol of each primer and 1 μL of a five-fold dilution of reverse transcriptase reaction product were used in a final reaction volume of 10 μL with PCR cycling conditions as follows: 9 min at 95°C, 45 cycles of 5 s at 95°C, 10 s at 65°C and 20 s at 72°C. At least three independent biological replicates, each ran in two technical replicates were performed. All reactions were checked for their dissociation curves. Relative expression was calculated using the ΔΔCp method with the *SAND* family gene At2g28390 as a constitutive control and analyzed with a type III ANOVA was used to analyze relative gene expressions. The post-hoc analysis consisted of multiple pairwise t-test. The adjusted p-values were obtained by Benjamini, Hochberg, and Yekutieli method (**25**) to control for the false discovery rate.

### Confocal imaging

MAP4-GFP Arabidopsis (**26**) leaves were imaged between 24 and 26 hours post-inoculation (hpi),hpi with infections by *S. sclerotiorum* expressing OAH1::GFP, using a Leica TCS TCSPC SP8 confocal microscope with a HC FLUOTAR 25x/0.95 water objective. Preparations were sealed with Vaseline to avoid evaporation and to limit spurious leaf displacements. Images were acquired at 1024×1024 pixels with 4x line average and a z-stack step of 0.8μm. GFP fluorescence was excited with the 488 nm ray line of an argon laser and recorded in the 505 to 530 nm emission range.

### Images analyses

Noise and isolated pixels in image stacks were removed by applying a median filter (Fiji > radius=3). The z-projection of the maximum pixel intensity was used as the input image for the computation of the CMT anisotropy, CMT angle and the displacement field (Fiji > Z Stack). A binary mask was drawn manually using a graphical tablet to separate the plasma membranes and cell walls from the cortical microtubules (CMTs). For every cell, the mean CMTs angle (in 0°, 180°) and anisotropy (0 = purely isotropic, 1= purely isotropic) were computed on masked images using fibriltool Fiji plugin (**27**). We used the ST+KLT method (**28**) of the open source Kineplant’s toolbox (https://sites.google.com/site/crtoolbox/home) to compute the spatial field of displacement (**29**). The maximal number of good features to track was set to 2000 and the size of the Gaussian interpolation window was set at 16 px. Cell walls and membrane displacements were obtained by convoluting the dense field of displacement given by the ST+KLT method and the inverted mask.

### Computation of inoculation-derived stretch directions and comparison with CMTs orientation

The displacement field of cell walls and plasma membranes obtained by the ST+KLT method was used for the computation of stretch directions. For every cell, a 2D linear transformation Φ was fitted to describe the spatial displacement of the cell wall over 2 hours. The transformation tensor **F** was computed as the gradient of Φ. Green-Lagrange strain tensor **E** was computed from **F** as **E** = (**F**^T^**F**-**I**)/2. The two principal components of the strain tensor **E** were deduced from its eigenvectors with eigenvalues λ_1_, λ_2_. Infection-derived stretch directions were compared to the mean direction of CMTs. For λ_1_, λ_2_ with opposite signs, the stretch direction was the direction of the eigenvector associated with the maximal positive eigenvalue. For negative λ_1_, λ_2_, no stretch direction was reported. For positive λ_1_, λ_2_, we assumed that CMTs reorganization occurred in both stretch directions. Consequently we computed the angle between the CMTs and the direction of eigenvectors bisector. The relative surface variation was computed as the sum of eigenvalues of **E**. Computations and stretch field rendering were performed using the NumPy python package (**30**) and matplotlib (**31**).

### Analyses of CMTs reorientation

The comparison of CMTs anisotropies in healthy and inoculated tissues was performed on 278 cells (109 in infected tissue, 169 in healthy tissue) corresponding to 6 different leaves (3 infected, 3 healthy) of 6 different *Arabidopsis* plants. Wilcoxon’s test was used to assess the mean difference in absolute values of the relative cell-surface variation for inoculated and healthy plants, and in phenotypic effect of impaired CMT dynamics on the susceptibility to *S. sclerotiorum* inoculation. The temporal evolution of CMT caused by pathogen-derived strain was tested by the linear model **φ**(t=26hpi) = a.**φ**(t=24hpi) +b+ε, with **φ** angles between the infection-derived stretch direction and the CMT direction. Slopes inferior to 1 indicated a convergence between CMTs angle and infection-derived stretches. We next tested variations in proportion within 30° angular sectors at constant time and between 24 hpi and 26 hpi by a proportion test, with results summarized by a letter in **Fig.1D, Fig2B,C**. All statistical analyses were performed using R software (**32**) and plots drawn using the ggplot2 library (**33**).

### Modeling mechanical stress and strains during infection

Healthy plant tissue was modeled as a flat membrane with a radius of 15 mm and thickness of 100 µm. The internal stress of plant tissue due to the interaction of water in vacuole and cell walls was modeled as an in-plane isotropic tension set to 100 N/m. The Young’s modulus of the membrane was set to 1 MPa and Poisson’s modulus set to 0.4. The principal stress associated with the initial tension was c.a. 0.3 MPa and in the same magnitude as reported turgor pressure. Fungal inoculation was modeled by reducing Young’s and Poisson’s modulus in-plane tension (**Table S2**) in a subdomain of the membrane. To test for spatial heterogeneities, the inoculated zone was modeled by two ellipses with a radius of 1 mm. Heterogeneities were modeled by mapping spatially different Young’s, Poisson’s, and in-plane tension. Parameter values were obtained by multiplying the value of the parameter in healthy tissue by a random number between 0.05 and 0.95. A sensitivity analysis was performed to test for the dependency of the overstretched area length on the lesion radius. Lesions were modeled as degraded tissue with lower Young’s modulus and lower in-plane tension. Different values of lesion radius, Young’s modulus, and in-plane tension were used to compute the stress field in healthy tissue (**Table S2**). Computation, modeling, and rendering were performed using COMSOL 5.2 software.

## Acknowledgments

The authors thank Dr. Bruno Moulia, Dr Olivier Hamant, Dr Stéphane Douady and Dr. Olivier Ali for helpful discussions regarding this work, Dr. Olivier Hamant and Dr. Marie-Edith Chabouté for providing the GFP expressing *Arabidopsis* seeds, the TRI-FR3450 Imaging Platform Facilities, Université de Toulouse, CNRS. The authors also thank the GDR Biophys and the labex TULIP (Grants ANR-10-LABX-41 and ANR-11-IDEX-0002-02) that allowed the formation of the interdisciplinary consortium gathered for this work.

