## Supplementary material for "Pathogen-derived mechanical cues regulate the spatio-temporal implementation of plant defense": Suplementary 1/1

**This PDF file includes:**

Figs. S1
Tables S1

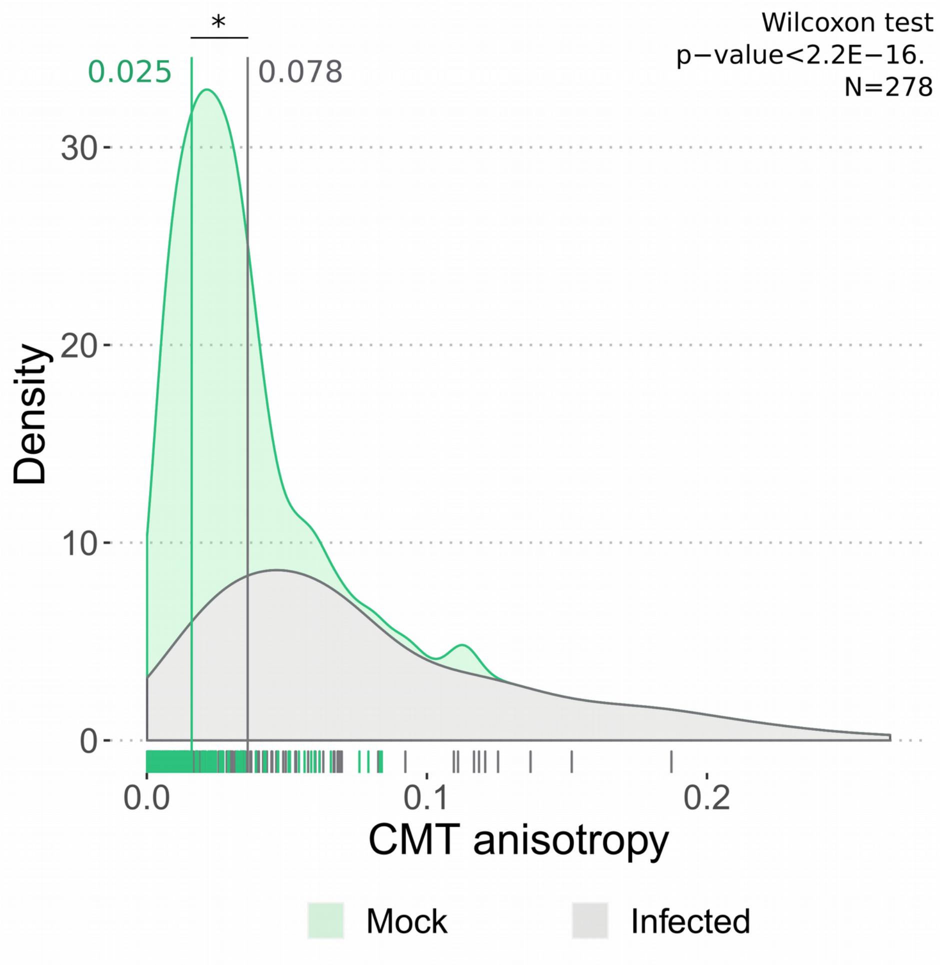

**Fig. S1.**

**Distributions of CMTs anisotropy in healthy and infected tissue.** Vertical bar indicates distributions means.

**Table S1**
Primer sequences used for quantitative RT-PCR.

| **Gene** | **Locus** | **Primer sequence** |
|---|---|---|
| AtMON1_Fw | At2g28390 | 5'- AACTCTATGCAGCATTTGATCCACT -3' |
| AtMON1_*Rev* | At2g28390 | 5'-TGATTGCATATCTTTATCGCCATC-3' |
| AtMCA1_Fw | AT4G35920 | 5'-TGCTCGGAACCTTCTCTTTGC-3' |
| AtMCA1_Rev | AT4G35920 | 5'-TTGGATGCTGCAGTGGCAATC-3' |
| AtBON1_Fw | AT5G61900 | 5'-AAGAAAAAGCGCGACTTAGACA-3' |
| AtBON1_Rev | AT5G61900 | 5'-GAGGAACCACTTCCACCAAC-3' |
| AtBON2_Fw | AT5G07300 | 5'-AATCCTGGCCTGGGAGATTC-3' |
| AtBON2_Rev | AT5G07300 | 5'-GAAAACCGCATGCGGGAAAG-3' |
| AtPDR12_Fw | AT1G15520 | 5'-TGAGCGAAAGCATAAGGCATGT-3' |
| AtPDR12_Rev | AT1G15520 | 5'-GGAGAAATCCTCCTTACACAGCC-3' |
| AtPRT_Fw | AT4G36010 | 5'-ACTTGTGGCGGAGCTGATTACG-3' |
| AtPRT_Rev | AT4G36010 | 5'-TCGTTGGACTCTTCACAGTTGGG-3' |

**Table S2.**

Parameters values used to model the effect of fungal-derived hydrolysis heterogeneities on the principal strain patterning and the dependency of the overstretched length on the lesion radius.

| Principal strain field patterning | | |
|---|---|---|
| | Healthy tissue | Lesion |
| Radius | 15 mm | 1 mm |
| Thickness | 0.1 mm | 0.1 mm |
| Young’s Modulus | 1 Mpa | In [0.05 , 0.95] MPa |
| Poisson's Modulus | 0.4 | In [0.02 , 0.38] |
| In-plane tension | 100 N/m | In [5 , 95] N/m |
| Overstretched length | | |
| Radius | 15 mm | In [1 , 10] mm |
| Thickness | 0.1 mm | 0.1 mm |
| Young’s Modulus | In {1, 5 , 10} MPa | In {0.01, 0.05, 0.1} Mpa |
| Poisson's Modulus | 0.4 | 0.2 |
| In-plane tension | In {10 , 100 , 1000} N/m | In {1 , 10 , 100} N/m |
